# Reproducible generation of human liver organoids (HLOs) on a pillar plate platform *via* microarray 3D bioprinting

**DOI:** 10.1101/2024.03.11.584478

**Authors:** Sunil Shrestha, Vinod Kumar Reddy Lekkala, Prabha Acharya, Soo-Yeon Kang, Manav Goud Vanga, Moo-Yeal Lee

## Abstract

Human liver organoids (HLOs) hold significant potential for recapitulating the architecture and function of liver tissues in vivo. However, conventional culture methods of HLOs, forming Matrigel domes in 6-/24-well plates, have technical limitations such as high cost and low throughput in organoid-based assays for predictive assessment of compounds in clinical and pharmacological lab settings. To address these issues, we have developed a unique microarray 3D bioprinting protocol of progenitor cells in biomimetic hydrogels on a pillar plate with sidewalls and slits, coupled with a clear bottom, 384-deep well plate for scale-up production of HLOs. Microarray 3D bioprinting, a droplet-based printing technology, was used to generate a large number of small organoids on the pillar plate for predictive hepatotoxicity assays. Foregut cells, differentiated from human iPSCs, were mixed with Matrigel and then printed on the pillar plate rapidly and uniformly, resulting in coefficient of variation (CV) values in the range of 15 - 18%, without any detrimental effect on cell viability. Despite utilizing 10 – 50-fold smaller cell culture volume compared to their counterparts in Matrigel domes in 6-/24-well plates, HLOs differentiated on the pillar plate exhibited similar morphology and superior function, potentially due to rapid diffusion of nutrients and oxygen at the small scale. Day 25 HLOs were robust and functional on the pillar plate in terms of their viability, albumin secretion, CYP3A4 activity, and drug toxicity testing, all with low CV values. From three independent trials of in situ assessment, the IC50 values calculated for sorafenib and tamoxifen were 6.2 ± 1.6 µM and 25.4 ± 8.3 µM, respectively. Therefore, our unique 3D bioprinting and miniature organoid culture on the pillar plate could be used for scale-up, reproducible generation of HLOs with minimal manual intervention for high-throughput assessment of compound hepatotoxicity.

## Introduction

Due to the urgent need for physiologically relevant *in vitro* tissue models for disease modeling and predictive compound screening, there have been rapid advancements made in the field of human organoids ^1^. With a better understanding of embryonic development, significant advances have been made in the differentiation of iPSCs into human liver organoids (HLOs) that can mimic the morphological features of human liver tissues and contain polarized hepatocytes with bile canaliculi-like architecture ^2,3^. In the earliest study of HLO development in 2013, the Takebe group generated liver buds by co-culturing pluripotent stem cell (PSC)-derived hepatic endoderm cells, human umbilical vein endothelial cells, and human mesenchymal stem cells, which were vascularized by transplantation in mice ^4^. Since then, several studies have been published with improved methods for the generation of HLOs through differentiation of iPSCs ^5–12^ or adult stem cells (ASCs) ^13^.

Despite notable achievements in generating physiologically relevant HLOs, several technical challenges remain for fully implementing HLOs for high-throughput screening (HTS) assays. These challenges include high costs associated with long-term cell differentiation, batch-to-batch variability, potential diffusion limitation of nutrients and oxygen into the core of large Matrigel domes, relatively low hepatic function of HLOs, and lack of high-throughput microfluidic devices for multi-organoid culture and communication ^14^. Conventionally, HLOs have been generated by either manually embedding foregut cells in Matrigel domes in 6-/24-well plates ^15^ or by the direct differentiation of PSCs seeded in basement membrane matrix-coated well plates and petri dishes^7^. However, these methods are labor-intensive, difficult to automate for HTS assays, and demand a large volume of cell culture media and reagents. To upscale organoid production, spinner flasks have been employed, improving the proliferation of organoids due to high oxygenation ^16^.

Additionally, a micropatterned cell-adhesion substrate (MPCS) in a 24-well plate was introduced by oxygen plasma treatment for high-throughput generation of functional HLOs ^17^. Despite improved organoid availability, these organoid culture systems are not easily amenable to HTS of compounds due to the need for isolation and encapsulation of HLOs in biomimetic hydrogels, imposing low throughput for downstream analyses. As a result, HLOs have been harvested typically during the late stage of differentiation from Matrigel domes through mechanical and enzymatic dissociation and then dispensed in high-density microtiter well plates, such as 384-well plates. The high-density microtiter well plates with hydrogel bottom coating could be used either for further differentiation and maturation followed by HTS assessment ^12^ or for direct, short-term (up to 72 hours) cell-based assays due to the instability of HLOs without hydrogel encapsulation^6^. While automation of HLO dispensing using robotic liquid dispensers appears feasible because of the relatively small diameter of HLOs (50 - 200 µm), the mechanical/enzymatic dissociation of HLOs in Matrigel domes could lead to the loss of organoids, damage to HLO structure and function, and batch-to-batch variation in their size.

Alternatively, microfluidic devices have been used to generate organoids, enhancing the maturity and physiological relevance of these organoids and allowing for *in situ* organoid-based assays ^18^. Nonetheless, most liver-on-chip systems developed so far have incorporated primary hepatocytes and non-parenchymal cells (NPCs) in microfluidic channels either to recapitulate specific hepatic features ^19–21^ or to enhance morphological and functional characteristics of liver constructs ^22–24^. These microfluidic devices are inherently low throughput and expensive to operate due to the requirement for tubes and syringe pumps ^25^. Recently, a perfusable chip system has been designed to generate multiple HLOs with improved hepatic functions ^26^, but it is low throughput in downstream analysis, requiring HLOs to be transferred to high-density well plates.

On the other hand, 3D bioprinting technology has been used either to create complex 3D cell models of tissue or to automate and upscale 3D culture systems for high-throughput assessment^27^. Several 3D liver models have been generated by 3D bioprinting, such as hepatic lobules using a mixture of liver cell types ^28–30^ and iPSC-derived hepatic models ^31,32^. Nonetheless, only a few studies have been performed for high-throughput culture of organoids by leveraging the automation capability of a 3D bioprinter such as an extrusion-based liquid handling system ^33^. Since iPSCs and organoids are highly fragile in nature, most extrusion-based 3D bioprinting systems would be unsuitable due to high shear stress, leading to relatively low cell viability (e.g., 40 - 86%) ^34^. To create bioprinted HLOs in high-throughput and reproducibly for compound screening, there are several critical criteria that need to be satisfied, including low shear stress in 3D bioprinting, nontoxic hydrogel gelation, a small size of bioprinted constructs to minimize the necrotic core and increase throughput, and compatibility with existing analytical equipment.

To address these requirements and achieve high-throughput organoid culture and *in situ* assessment of organoids with compounds, new engineering approaches are urgently necessary. In the present work, we developed a pillar plate with sidewalls and slits by injection molding of polystyrene and demonstrated reproducible, miniature, and scalable HLO culture on the pillar plate *via* microarray bioprinting of foregut cells in Matrigel for *in situ* assessment of compound hepatotoxicity. Scalability in culture, reproducibility in organoid differentiation, maturity of organoids generated, and high throughput in organoid-based assays are critical features necessary for HTS of compound libraries in pharmaceutical industries ^35^.

## Materials and Methods

### Fabrication of pillar and deep well plates

A 36-pillar plate with sidewalls and slits (36PillarPlate) and a 384-pillar plate with sidewalls and slits (384PillarPlate) were manufactured by the injection molding of polystyrene (**Figure 1**) and functionalized with poly(maleic anhydride-*alt*-1-octadecene) (PMA-OD; Sigma, 419117) and alginate (Sigma; A1112) for cell printing and organoid culture (Bioprinting Laboratories Inc., Dallas, TX, USA) ^36,37^. The 36PillarPlate and the 384PilllarPlate each contain a 6 x 6 array of pillars and a 16 x 24 array of pillars (4.5 mm pillar-to-pillar distance, 11.6 mm pillar height, and 2.5 mm outer and 1.5 mm inner diameter of pillars), respectively (**Supplementary Figure 1**). For organoid culture, the pillar plate with cells encapsulated in Matrigel was inverted and sandwiched onto a clear-bottom 384-deep well plate (384DeepWellPlate) manufactured by the injection molding of polystyrene (Bioprinting Laboratories Inc.). The 384DeepWellPlate has a 16 x 24 array of deep wells (3.5, 3.5, and 14.7 mm well width, length and depth, and 4.5 mm well-to-well distance) for cell culture (**Supplementary Figure 2**). The unique structure of sidewalls and slits on the pillars and alginate coating on the surface prevent detachment of cell-containing Matrigel droplets from the pillar plate and 2D growth of cells on the surface during long-term organoid culture.

**Figure 1.**
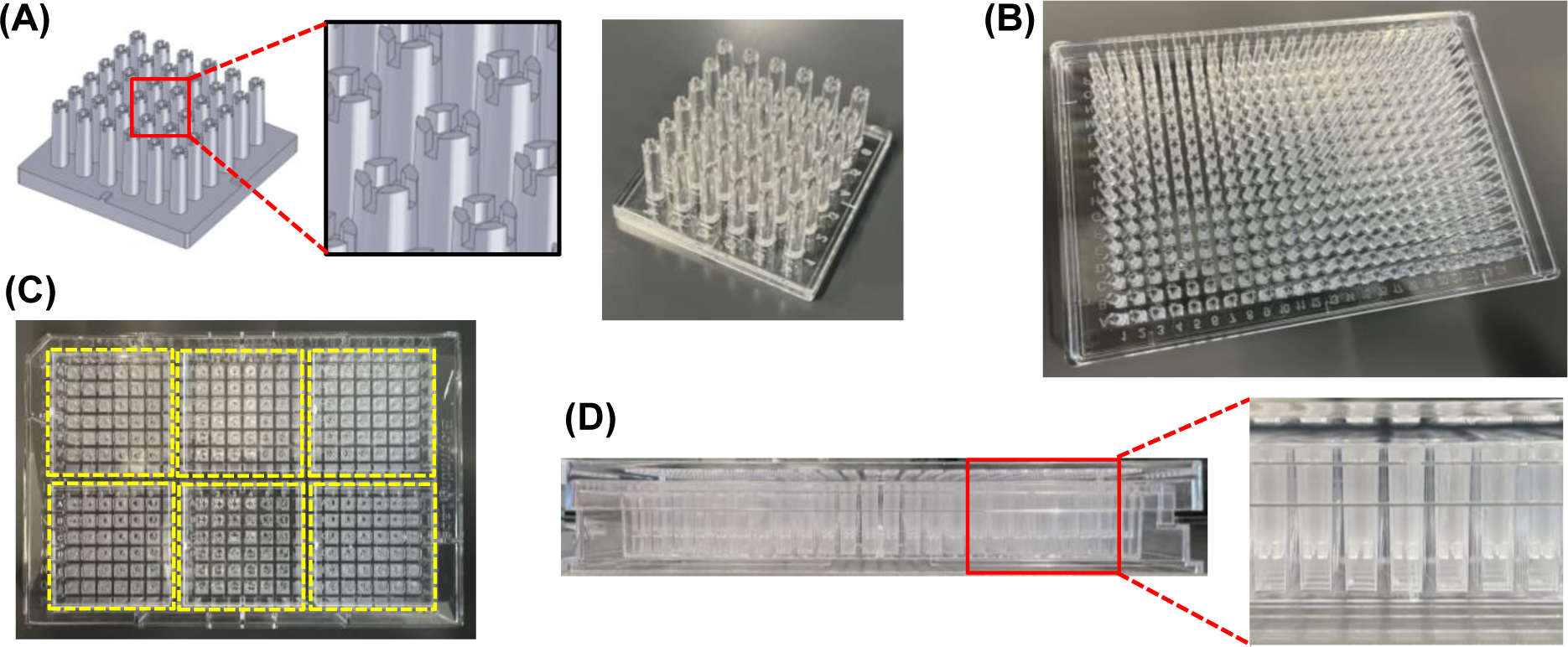
Injection molding of a pillar plate and a complementary deep well plate for static organoid culture: **(A)** SolidWorks design of a 36PillarPlate with a 6 x 6 array of pillars and the picture of an injection-molded 36PillarPlate. **(B)** An injection-molded 384PillarPlate with a 16 x 24 array of pillars. **(C)** Top view of six 36PillarPlates sandwiched onto a 384DeepWellPlate for static organoid culture. **(D)** Side view of the 384PillarPlate sandwiched onto the 384DeepWellPlate.

### Maintenance and differentiation of iPSCs into foregut cells

EDi029-A, a male human iPSC line (Cedar Sinai Biomanufacturing Center, USA), was maintained on growth factor reduced Matrigel (Corning; 354230) coated dishes using mTeSR^TM^ Plus medium (Stemcell Technologies; 100-0276). When healthy colonies of cells were formed, they were passaged to form uniform cell clusters using the StemPro^TM^ EZPassage^TM^ passaging tool (ThermoFisher; 23181-010). The differentiation of iPSCs into foregut cells was performed using previously published protocols ^6,12^. Briefly, at 70 - 80% confluency, iPSCs were harvested using Accutase (Gibco; A1110501), seeded on iMatrix-511 silk (Elixirgen Scientific; NI511) coated 6-well plate at a cell density of 1.3 x 10^6^, and cultured using mTESR^TM^ Plus medium supplemented with 10 µM Y27632 Rho-kinase (ROCK) inhibitor (Tocris; 1254). After 24 hours of culture, the iPSCs were differentiated into definitive endoderm using RPMI 1640 (Gibco; 22400089) supplemented with 50 ng/mL bone morphogenetic protein 4 (BMP4; Tocris; 314-BP) and 100 ng/mL activin A (Tocris; 338-AC) at day 1, 100 ng/mL Activin A and 0.2% knockout serum replacement (KSR; Gibco; 10828010) at day 2, and 100 ng/mL Activin A and 2% KSR at day 3. This was followed by foregut cell differentiation from day 4 - 6 using advanced DMEM/F12 (Gibco; 12634) with 2% B27 (Gibco; 17504), 1% N2 (Gibco; 17502), 10 mM HEPES (Gibco; 15630), 1% Pen/Strep (Gibco; 15140), and GlutaMAX^TM^ (Gibco; 35050) supplemented with 500 ng/mL fibroblast growth factor (FGF4; Peprotech; 100-31) and 3 mM CHIR99021 (R&D Systems; 4423).

### Microarray bioprinting of cells in Matrigel on the pillar plate

The suspension of cells in Matrigel were printed using ASFA^TM^ Spotter (MBD Korea; V6) (**Supplementary Figure 3**). Before printing, iPSCs or day 7 foregut cells were suspended in undiluted Matrigel (Corning; 356237). The critical parameters such as open time (µs) and pressure (kPa) were set using medical and bio-decision (MBD) software, which determines the printing volume per droplet (or spot) (**Supplementary Figure 4**). The pillar plate can accommodate up to 6 µL of cell-containing hydrogels at the tip without overflow, but we printed 4 - 5 µL of cell-Matrigel spots. The printing area on the pillar plate was selected using the software, which automatically calculates the volume of cell suspension in Matrigel necessary in the disposable printing tip with a 400 µm orifice for successful printing with minimal dead volume. For printing the entire 36PillarPlate with a 5 µL spot per pillar, it requires at least 180 µL of cell suspension in Matrigel. The pillar plate was placed in the position of the ‘target plate’ within the 3D bioprinter (**Supplementary Figure 3**). The necessary volume of cell suspension in Matrigel in the printing tip can be changed depending on the dispensing volume and the number of the pillar plates printed. To print the cell suspension in Matrigel on the entire 36PillarPlate and the 384PillarPlate, it takes approximately 20 and 60 seconds, respectively. There was no precipitation of cells in hydrogel on the pillar plate during printing due to the extremely fast printing speed.

### Differentiation of foregut cells into HLOs in Matrigel on the pillar plate

On day 7, foregut cells dissociated with Accutase were either cryopreserved using CryoStor^®^ CS10 (Stemcell technologies; 07959) for later use or mixed with Matrigel (Corning; 356237) at a density of 750 cells/µL for culture. The cell suspension in Matrigel was either dispensed in a 24-well plate to form 50 µL domes or printed on the pillar plate at a 4 µL spot per pillar. After gelation of Matrigel at 37°C for 10 - 12 minutes, foregut cells in the 24-well plate or on the pillar plate were cultured in advanced DMEM/F12 supplemented with 5 ng/mL recombinant human FGF basic/FGF2/bFGF (R&D systems; 233-FB), 10 ng/mL recombinant human VEGF-165 (Gibco; PHC9391), 20 ng/mL recombinant human EGF (R&D system; 236-EG), 0.5 µM A 83-01 (R&D Systems; 2939), 50 µg/mL L-ascorbic acid (Sigma; A4544), and the CEPT cocktail consisting of 50 nM chroman 1 (R&D systems; 7163), 5 µM emricasan (Selleckchem; S7775), 1x polyamine supplement (Sigma; P8482), and 0.7 µM trans-ISRIB (R&D systems; 5284). The medium volume used in the 24-well plate was 1 mL per well. The pillar plate with bioprinted foregut cells was sandwiched with the complementary 384DeepWellPlate containing 80 µL culture medium in each well. The differentiation medium was changed every other day. On day 11, the differentiation medium was changed to advanced DMEM/F12 supplemented with 2 µM retinoic acid (Sigma; R2625) with medium change on every other day. From day 15 until day 25, the maturation medium consisting of HCM^TM^ Hepatocyte Culture Medium BulletKit^TM^ (Lonza; CC-3198), except no EGF supplied from the manufacture, supplemented with 10 ng/mL recombinant human hepatocyte growth factor (HGF; Peprotech; 100-39), 20 ng/mL recombinant human oncostatin M (OSM; Peprotech; 300-10), and 100 nM dexamethasone (DEX; Sigma; D4902) was used for HLO maturation and replaced every other day.

### Measurement of cell viability

The viability of cells was analyzed by using the CellTiter-Glo^®^ 3D cell viability assay kit (Promega; G9681) following the manufacturer’s recommended protocol. Briefly, the pillar plate with HLOs was sandwiched with an opaque white 384-well plate containing a mixture of 30 µL of the CellTiter-Glo^®^ reagent and 10 µL of the cell culture medium in each well to measure cellular adenosine triphosphate (ATP) levels. To induce cell lysis, the sandwiched pillar/well plates were placed on an orbital shaker for 1 hour at room temperature. Later, the pillar plate was detached, and the lysis solution in the opaque white 384-well plate was left for 15 minutes at room temperature for stabilization. The luminescence signals were recorded using a microtiter well plate reader (BioTek^®^ Cytation 5).

To measure the viability of bioprinted cells on the pillar plate, staining with calcein AM and ethidium homodimer-1 was performed. Briefly, the cells on the pillar plate were rinsed once with a saline solution and then stained with 80 µL of a mixture containing 2 µM calcein AM and 4 µM ethidium homodimer-1 in a 384DeepWellPlate for 1 hour at room temperature. After staining, the pillar plate was rinsed twice with the saline solution, and fluorescent images were acquired in high throughput with the Keyence BZ-X710 automated fluorescence microscope (Keyence, Osaka, Japan) at an excitation/emission of 494/517 nm for calcein AM and 528/617 nm for ethidium homodimer.

### Gene expression analysis *via* qPCR

Monolayer-cultured cells were collected *via* Accutase dissociation. HLOs in Matrigel were either collected manually through pipetting in cold PBS^-/-^ or isolated from Matrigel using Cultrex organoid harvesting solution (R&D Systems; 3700-100-01) according to the manufacturer’s recommended protocol, which allows non-enzymatic depolymerization of Matrigel. In case of HLOs on the pillar plate, the pillar plate was sandwiched onto the deep well plate containing 80 µL of Cultrex organoid harvesting solution. The sandwiched plates were incubated for 30 minutes at 4°C and then centrifuged at 100 rcf for 10 minutes to detach the organoids. Total RNA was isolated from harvested cells by using the RNeasy Plus Mini Kit (Qiagen; 74134) following the manufacturer’s recommended protocol. cDNA was synthesized from 1 µg of RNA by following the protocol of the high-capacity cDNA reverse transcription kit (Applied Biosystems; 4368814). Real time PCR was performed using PowerTrack^TM^ SYBR green master mix (Applied Biosystems; A46110) and forward/reverse primers from IDT Technology in the QuantStudio™ 5 Real-Time PCR System (Applied Biosystems; A28574). The cycle was run 40 times at 95°C denaturation for 30 sec, 58 - 62°C annealing for 45 sec, depending on primer pair, and 72°C extension for 30 sec. The primers used are listed in **Supplementary Table 1**. The expression level of target genes was normalized with that of the housekeeping gene, glyceraldehyde 3-phosphate dehydrogenase (*GAPDH*).

### Whole mount immunofluorescence staining of HLOs

The immunofluorescence staining was performed either by harvesting HLOs from Matrigel domes in the 24-well plate or with HLOs *in situ* on the pillar plate. In case of Matrigel dome culture, Matrigel domes containing organoids were collected in cold dPBS^-/-^ through pipetting into a 1.5 mL Eppendorf tube and then centrifuged to isolate HLOs at 300 x g for 4 minutes. The HLOs were fixed using 4% paraformaldehyde (PFA; Thermo Scientific; J19943K2) for 2 hours at room temperature while gently rocking. The fixed HLOs were washed with 0.1% (w/v) sodium borohydride in dPBS^-/-^ twice for 15 minutes to reduce background due to free aldehyde. After washing, the HLOs were permeabilized with 500 µL of 0.5% Triton X-100 (Sigma; T8787) in dPBS^-/-^ (i.e., permeabilization buffer) for 15 - 20 minutes twice at room temperature with gentle rocking. After permeabilization, HLOs were exposed to 500 µL of 5% normal donkey serum (NDS) in the permeabilization buffer (i.e., blocking buffer) for 1 hour at room temperature or overnight at 4°C with gentle rocking to prevent non-specific binding. For primary antibody staining, the HLOs were treated with 250 µL of a 5 µg/mL primary antibody diluted in the blocking buffer for overnight at 4°C with gentle rocking. The HLOs were rinsed with 1 mL of the blocking buffer thrice for 30 minutes each at room temperature with gentle rocking to prevent non-specific binding. For secondary antibody staining, the HLOs were exposed to 500 µL of 5 µg/mL fluorophore-conjugated secondary antibody in the blocking buffer for 2 - 4 hours at room temperature with gentle rocking. The HLOs were stained with 500 µL of 0.5 µg/mL DAPI solution (ThermoFisher Scientific; 62248) in 1x dPBS^-/-^ for 30 minutes at room temperature with gentle rocking. The HLOs were further washed with 1 mL of dPBS^-/-^ twice to ensure the complete removal of unbound secondary antibody. Finally, the HLOs were transferred to a microscope cover glass (FisherScientific; 22266882) and treated with 25 μL of Visikol^®^ Histo-M™ (Visikol; HM-30) to clear the organoids which also works as a mounting solution. The HLOs on the cover glass slide were covered by another cover glass from the top and imaged using a Zeiss LSM 710 confocal scanning microscope. For the HLOs on the pillar plate, all the immunofluorescence staining steps were performed by sandwiching the pillar plate with a 384DeepWellPlate containing 80 µL of respective solutions and incubating the sandwiched plates under the same conditions mentioned above. After the final wash with 80 μL dPBS^-/-^ to remove unbound secondary antibody, the HLOs were treated with 35 µL of Visikol^®^ Histo-M™ in a regular 384-well plate (Thermofisher Scientific; 242757) for 1 hour at room temperature. At the time of imaging, the pillar plate containing stained HLOs was placed on the microscope cover glass. The specific names of primary and secondary antibodies used are listed in **Supplementary Table 2 and 3**, respectively.

### Cell cytometry analysis

Foregut cells were dissociated into single cells with Accutase for 5 - 8 minutes at 37°C inside a CO2 incubator. The dissociated cells were collected in a 15 mL tube and fixed using 4% PFA for 15 minutes at room temperature. Subsequently, the cells were permeabilized with 0.25% Triton X-100 in PBS for 15 minutes and blocked using 5% NDS in the permeabilization solution for 30 minutes at room temperature. The cells were then incubated with a 5 µg/mL of human HNF-3 beta/FoxA2 antibody (R&D Systems; AF2400) for 1.5 hours at room temperature, washed with the blocking solution thrice, and stained with a 5 µg/mL of donkey anti-goat IgG NorthernLights™ NL637-conjugated antibody (R&D Systems; NL002) for 30 minutes at room temperature. Subsequent to staining, the cells were washed with PBS three times before flow cytometry analysis. UltraComp eBeads™ compensation beads (Fisher Scientific; 01-2222-41) stained with the same secondary antibody were used as a positive control for analysis. The analysis was performed by the Cytek Aurora spectral flow cytometer (Cytek^®^ Biosciences) and Floreada website (https://floreada.io/).

### Measurement of albumin secretion with an ELISA assay

To measure the level of albumin secretion from HLOs on the pillar plate, 80 μL of the culture medium in the 384DeepWellPlate was collected at day 25 of culture after 48 hours of incubation with HLOs encapsulated in Matrigel on the pillar plate. The collected culture medium was centrifuged at 250 g for 3 minutes to remove any debris, and the resulting supernatant was assayed using a human albumin ELISA kit (ThermoFisher Scientific; EHALB) according to the manufacturer’s instruction.

### Measurement of CYP3A4 expression

The expression level of CYP3A4 was analyzed by using the P450-Glo^TM^ CYP3A4 assay kit (Promega; V9001) and following the manufacturer’s recommended protocol. Rifampicin (Sigma; R3501) was used as an inducer of the CYP3A4 gene at the concentration of 25 µM. Briefly, on day 25, HLOs on the pillar plate were treated with rifampicin for 3 days with daily medium changes. After treatment, HLOs were incubated with luciferin IPA-substrate at a final concentration of 3 µM diluted in basal HCM medium (without EGF), overnight at 37°C in the CO_2_ incubator. Following the overnight incubation, 25 µL of the culture medium from each well of the 384DeepWellPlate was transferred to the opaque white 384-well plate at room temperature, and 25 µL of luciferin detection reagent was added in each well to initiate a luminescent reaction. After 20 minutes of incubation at room temperature, luminescence was measured by using the BioTek^®^ Cytation 5 plate reader.

### Measurement of bile acid transport with cholyl-lysyl-fluorescein (CLF)

To analyze the function of bile acid transport in HLO, HLOs formed in 50 µL Matrigel domes were treated with 5 µM cholyl-lysyl-fluorescein (AAT Bioquest Inc., CA, USA) in complete maturation medium. After overnight treatment at 37°C in the CO_2_ incubator, the HLOs were rinsed with dPBS^-/-^ thrice. The HLOs were then imaged by using the Zeiss LSM 710 confocal scanning microscope equipped with a 40x water immersion objective lens.

### Testing model compounds with HLOs on the pillar plate

Day 25 HLOs on the pillar plate were exposed to varying concentrations of two hepatotoxic drugs, including sorafenib and tamoxifen. The highest dosage tested was 200 μM for sorafenib and 500 μM for tamoxifen. Briefly, 4-fold serial dilutions of the highest dose of the drugs were performed in DMSO in a 384-well plate. Five dosages and one solvent-alone control (DMSO control) were prepared for each drug. The drug stock solutions in DMSO in the 384-well plate were 200-fold diluted with the maturation medium and then dispensed in the 384DeepWellPlate (six replicates per dose) so that the final DMSO content is equal to 0.5% (v/v). The pillar plate with day 25 HLOs was then sandwiched with the 384DeepWellPlate containing the serially diluted drugs and incubated in the 5% CO_2_ incubator at 37°C for 3 days. The viability of HLOs was assessed with CellTiter-Glo^®^ 3D cell viability assay kit (Promega), and luminescence was measured by using the BioTek^®^ Cytation 5 plate reader. Dose-response curves were generated using the luminescence values at varying dosages.

### Calculation of the IC_50_ value

Since the background luminescence of completely dead cells (following treatment with 70% methanol for 1 hour) was negligible due to background subtraction, the percentage of live HLOs was calculated using the following equation:

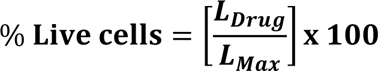

where L_Drug_ is the luminescence intensity of HLOs exposed to the drugs and L_Max_ is the luminescence intensity of fully viable HLOs (control).

To generate a conventional sigmoidal dose-response curve with response values normalized to span the range from 0% to 100% plotted against the logarithm of test concentration, we normalized the luminescence intensities of all HLO spots with the luminescence intensity of a 100% live HLO spot (HLOs incubated with no compound). We then converted the test drug concentrations to their respective logarithms using GraphPad Prism 9.3.1 (GraphPad Software, Inc., CA, USA). The sigmoidal dose-response curve (variable slope) and IC_50_ value (i.e., the concentration of drug where 50% of HLO viability is inhibited) were obtained using the following equation:

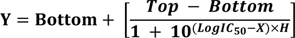

where IC_50_ is the midpoint of the curve, H is the hill slope, X is the logarithm of test concentration, and Y is the response (% live cells), starting from the top plateau (Top) of the sigmoidal curve to the bottom plateau (Bottom).

### Calculation of the coefficient of variation (CV)

To establish the robustness of cell printing on the pillar plate, the range of errors was measured using the coefficient of variation (CV). The CV value is calculated as the ratio of the standard deviation (SD) to the average (Avg). It serves as a measure of variability in relation to the average signal intensity, essentially representing the inverse of the signal-to-noise ratio.

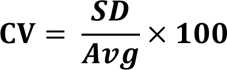

Thus, a low CV value is always preferred. In general, the acceptable range of CV values for cell-based assays in microtiter well plates are between 20 – 30 whereas a CV value greater than 30 is unacceptable ^38^.

### Statistical analysis

The statistical analysis was performed using GraphPad Prism 9.3.1. All the data were expressed as mean ± SD, with sample sizes specified in the figure legends where ‘n’ represents biological replicates. Student’s t-test was used for comparison between two groups, whereas one-way ANOVA was used for comparison among multiple groups. The statistically significant difference between the control and test groups was indicated by *** for p < 0.001, ** for p < 0.01, * for p < 0.05, and ns = not significant (p > 0.05).

## Results

### Uniform printing of iPSCs and foregut cells on the pillar plate using a 3D bioprinter

For scale-up production of human organoids and high-throughput organoid-based compound screening, it is imperative to automate the process of cell loading in the cell culture system and reduce the assay volume by miniaturization. Due to recent advances in 3D bioprinting, cells in hydrogels can be dispensed uniformly by several bioprinting techniques ^39^. However, the viability of bioprinted cells can be influenced strongly by gelation mechanisms and shear stress applied to the cells ^33^. In the present work, we employed a microsolenoid valve-driven, 3D bioprinting technique to print iPSCs and foregut cells suspended in thermosensitive hydrogels such as Matrigel on the pillar plate (**Figure 2A and Supplementary Figure 3**). This printing method allowed the dispensing of Matrigel in the range of 100 nL – 6 µL per spot, depending on the valve open time. In addition, it took approximately 20 and 60 seconds to dispense cells suspended in Matrigel on the entire 36PillarPlate and 384PillarPlate, respectively **(Figures 1 and 2)**. After printing cell suspension in Matrigel on the pillar plate, thermal gelation at 37°C for 10 - 12 minutes was performed. The viability of bioprinted iPSCs and foregut cells on the pillar plate was very high and almost identical to its non-printed control (**Figure 2**), which could be due to the low shear stress applied during cell printing and nontoxic thermal gelation of the hydrogels. The live/dead staining with calcein AM and ethidium homodimer-1 indicated high viability of bioprinted iPSCs in Matrigel on the pillar plate across three independent trials (**Figure 2B**). Although iPSCs are highly fragile, the viability of bioprinted iPSCs was identical to the viability of manually dispensed iPSCs with a pipette, which was measured by an ATP-based luminescence assay (**Figure 2C**). These results indicate that our microarray 3D bioprinting has no detrimental effect on cell viability and can support rapid cell loading on the pillar plate. In addition, the robustness of cell printing has been demonstrated by measuring the coefficient of variation (CV) with bioprinted foregut cells in Matrigel on the pillar plate (**Figure 2D**). The CV values obtained from the three trials were 16%, 15%, and 18%, indicating the robustness of the bioprinting approach and the high reproducibility of cell printing among the pillar plates.

**Figure 2.**
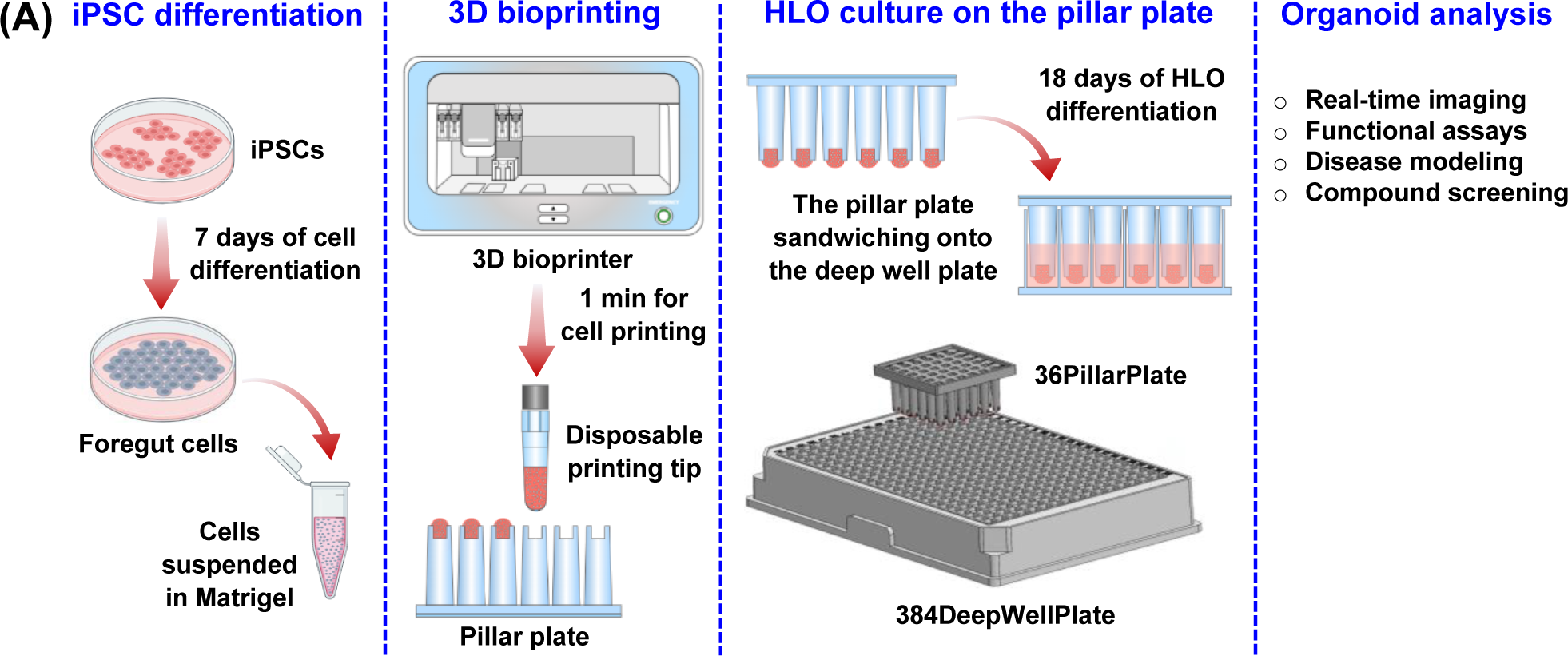
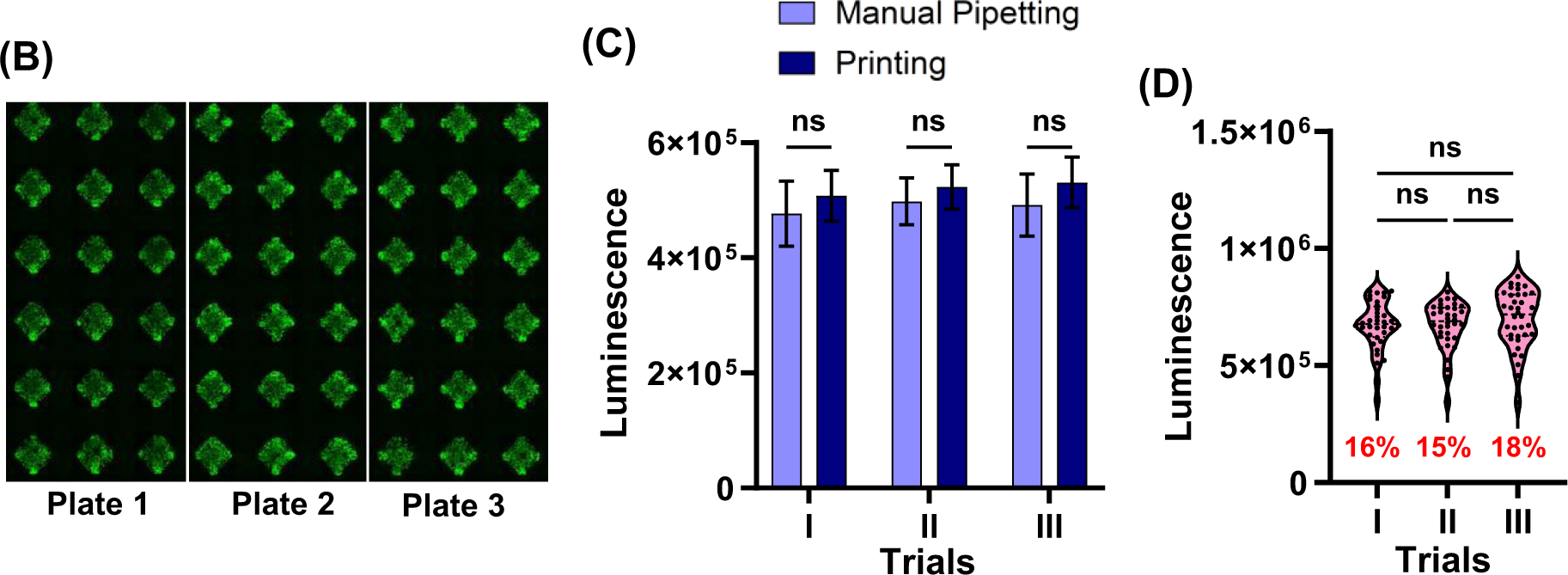
High-throughput printing of human cells in Matrigel on a pillar plate: **(A)** The process of iPSC differentiation into foregut cells, bioprinting of foregut cells in Matrigel on the pillar plate, and foregut cell differentiation into HLO for *in situ* organoid analysis. **(B)** Cell viability after printing iPSCs in Matrigel measured with calcein AM and ethidium homodimer-1 staining. Comparison of cell viability between bioprinted and manually dispensed iPSCs in Matrigel on the pillar plate by an ATP-based luminescence assay. The data were shown as mean ± SEM. n = 36. **(D)** Viability of bioprinted foregut cells in Matrigel on the pillar plate by the ATP-based luminescence assay. n = 36.

### Differentiation of iPSCs into HLOs in Matrigel domes in the 24-well plate

Before generating HLOs on the pillar plate, a male iPSC line from Cedars Sinai (EDi029-A) was differentiated into HLOs in Matrigel domes in a 24-well plate by following the protocol established by Ouchi et al. and Shinozawa et al. ^6,12^ (**Figure 3A**). Among several HLO differentiation protocols published, the protocol developed by the Takebe group allowed the generation of multicellular HLOs with higher liver gene regulatory network (GRN) and classification scores assessed by CellNet analysis ^40^. On day 7 of differentiation, the progenitor cells expressed representative foregut cell markers such as FOXA2 and SOX2, attesting to the foregut stage (**Figure 3B**). In addition, cell cytometry analysis showed a 96% FOXA2+ cell population, indicating high efficiency of iPSC differentiation into foregut cells (**Figure 3C**). The foregut cells were suspended in Matrigel and seeded in 50 µL Matrigel domes in the 24-well plate. Early hepatic progenitor cells were formed on day 15 after retinoic acid (RA) treatment and further differentiated up to day 25 in the maturation medium. The stepwise differentiation of iPSCs into HLOs was assessed by RT-qPCR analysis using specific hepatic markers for different stages of cell differentiation. The heatmap generated from the RT-qPCR analysis showed the iPSCs differentiated into the hepatic lineage by expressing early hepatic progenitor markers such as HNF4a and AFP and expressing ALB at the stage of mature HLOs (**Figure 3D**). In addition, whole-mount immunofluorescence staining showed the expression of albumin marker ALB, epithelial marker E-cad, hepatocyte marker HNF4a, and mesenchymal origin stellate cell marker VM (**Figure 3E**). Furthermore, the accumulation of green, fluorescent bile acid CLF at the intra-lumen of an HLO indicates the presence of an efflux transporter BSEP and bile acid transport activity in the HLO (**Figure 3F**).

**Figure 3.**
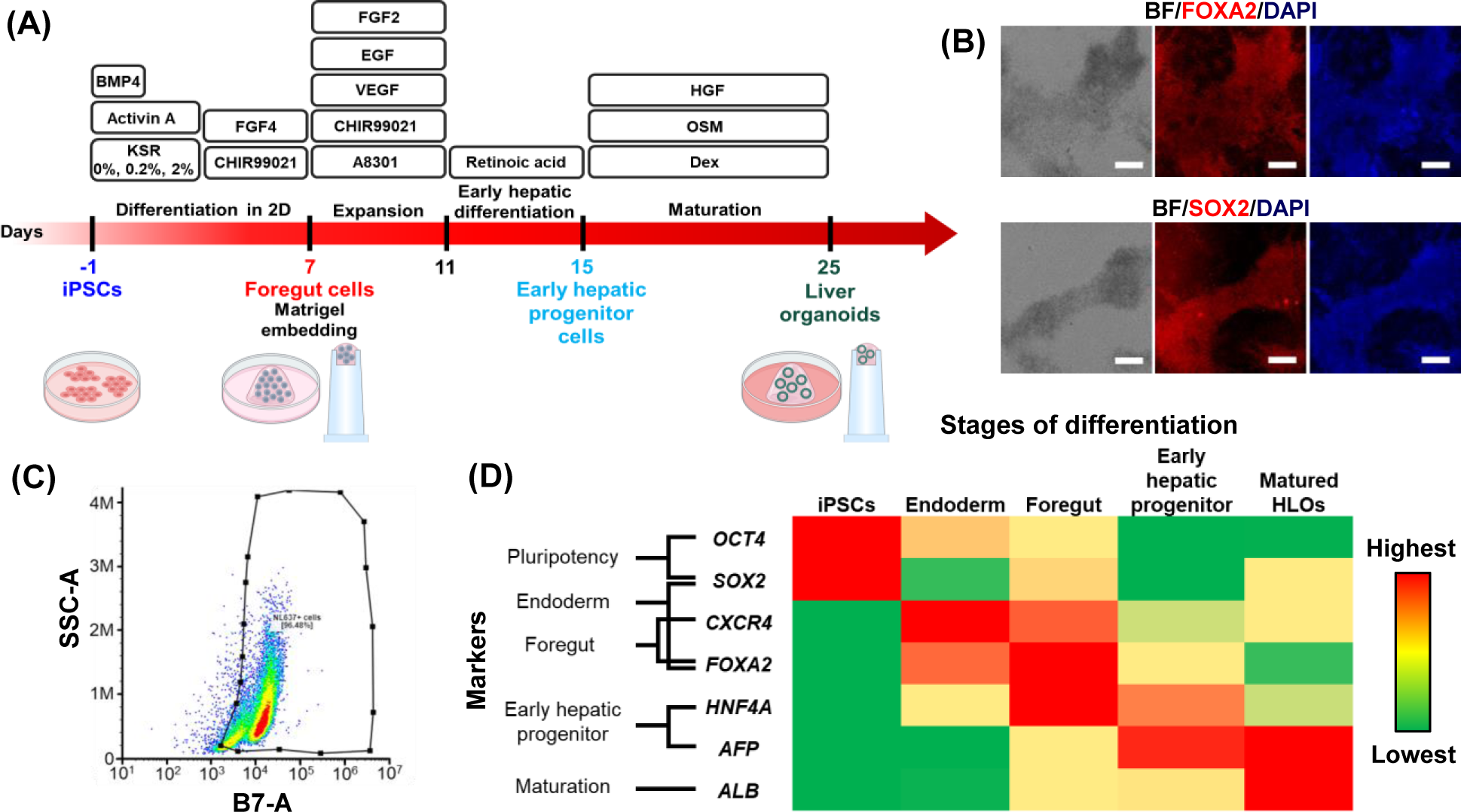
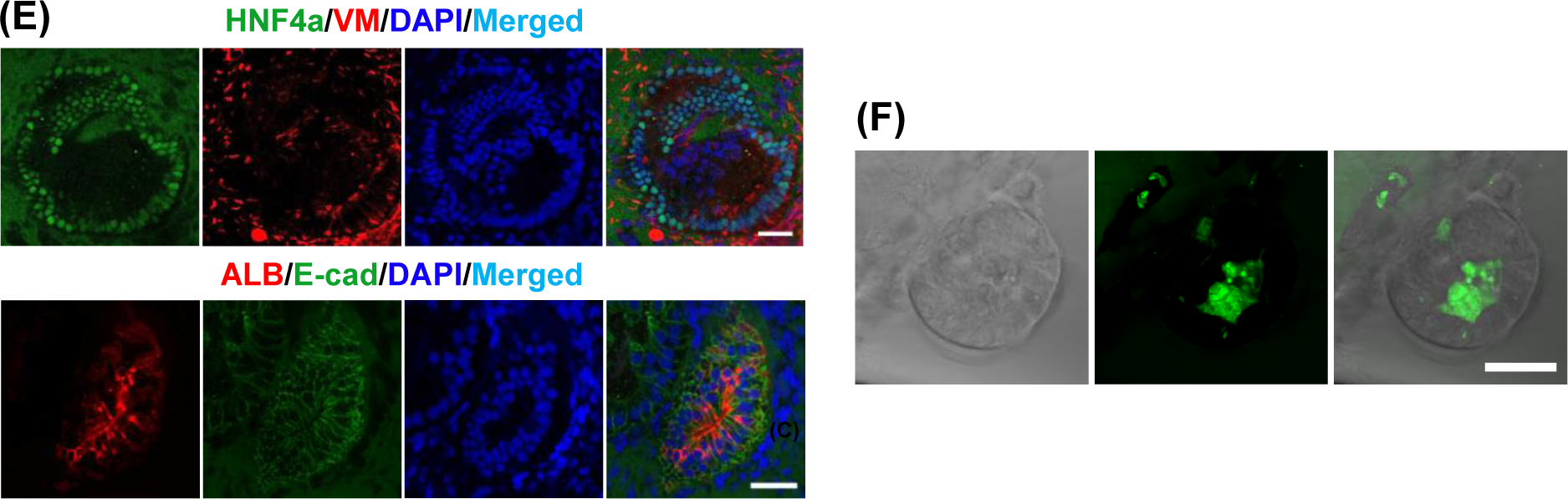
Generation of functional HLOs from EDi029A iPSCs. **(A)** HLO differentiation protocol in a 24-well plate and on the pillar plate. **(B)** Immunofluorescence staining of day 7 foregut cells for FOXA2 and SOX2 markers. Scale bars: 200 µm. **(C)** Scattered plot showing 96% FOXA2+ population in day 7 foregut cells. **(D)** Heatmap denoting the identity of cell population at different stages of differentiation generated based on relative gene expression by RT-qPCR analysis. n = 4. **(E)** Immunofluorescence staining of day 25 HLOs generated in 50 µL Matrigel dome culture. Scale bars: 50 µm. **(F)** Uptake of fluorescein-labeled bile acid, chilly-lysyl-fluorescein (CLF). Scale bar: 50 µm.

### Reproducible differentiation of iPSCs into HLOs in Matrigel spots on the pillar plate

Miniaturized organoid culture has been demonstrated by differentiating foregut cells in Matrigel on the pillar plate for 25 days (**Figure 4A**). In comparison to the conventional 50 µL Matrigel dome culture of HLOs in the 24-well plate, we demonstrated the feasibility of generating HLOs on the pillar plate by printing 4 µL of foregut cells suspended in Matrigel using the 3D bioprinter. The bioprinted foregut cells were differentiated into HLOs on the pillar plate by following the protocol developed by the Takebe group (**Figure 3A**). The relative gene expression levels of representative hepatic biomarkers, including *ALB* albumin, *ASGR1* hepatocytes, *SOX9* cholangiocytes, *VM* stellate cells, and *CD68* Kupffer cells, were either similar or higher in HLOs cultured on the pillar plate compared to those generated in conventional Matrigel dome culture (**Figure 4B**). In addition, the relative expression levels of *HNF4A* early hepatic progenitor cells, drug metabolizing enzymes such as *CYP3A4*, *UGT1A1*, *SULT2A1*, and *F7* coagulation factor VII were comparable between the HLOs cultured on the pillar plate and those cultured in the 24-well plate (**Figure 4B**). Furthermore, the secretion level of albumin from HLOs on the pillar plate was approximately 2-fold higher than that from the conventional Matrigel dome culture (**Figure 4C**). The enhancement in gene expression and albumin secretion may be attributed to the fast diffusion of nutrients and oxygen, facilitated by the 12-fold smaller cell culture on the pillar plate. Lastly, whole-mount immunofluorescence staining of HLOs on the pillar plate showed the expression of ALB, E-cad, HNF4a, and VM comparable to those in HLOs generated by conventional Matrigel dome culture (**Figure 4D**). The immunofluorescence staining of whole HLOs on the pillar plate showed multiple functional organoids surrounded by mesenchymal cells (**Figure 4E**). We have also generated day 25 HLOs on the pillar plate using foregut cells from patient-derived 72-3 iPSCs provided by our collaborator, the Takebe group at Cincinnati Children’s Hospital Medical Center, OH, USA (**Supplementary Figure 5**).

**Figure 4.**
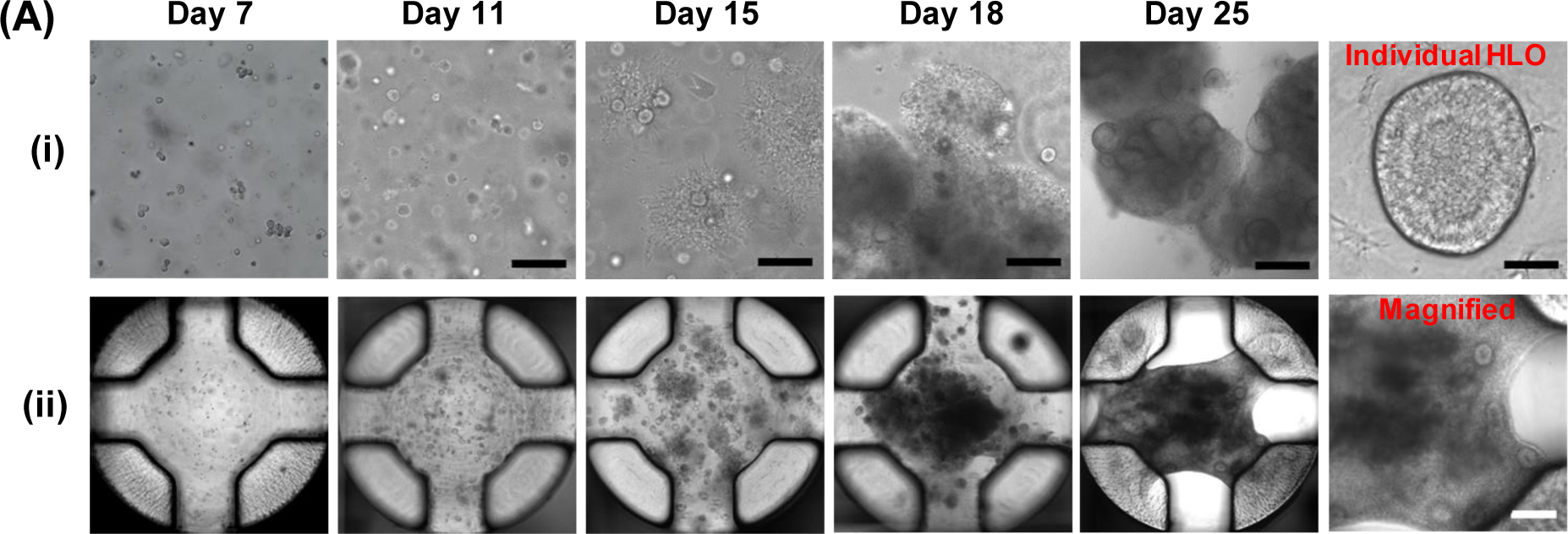
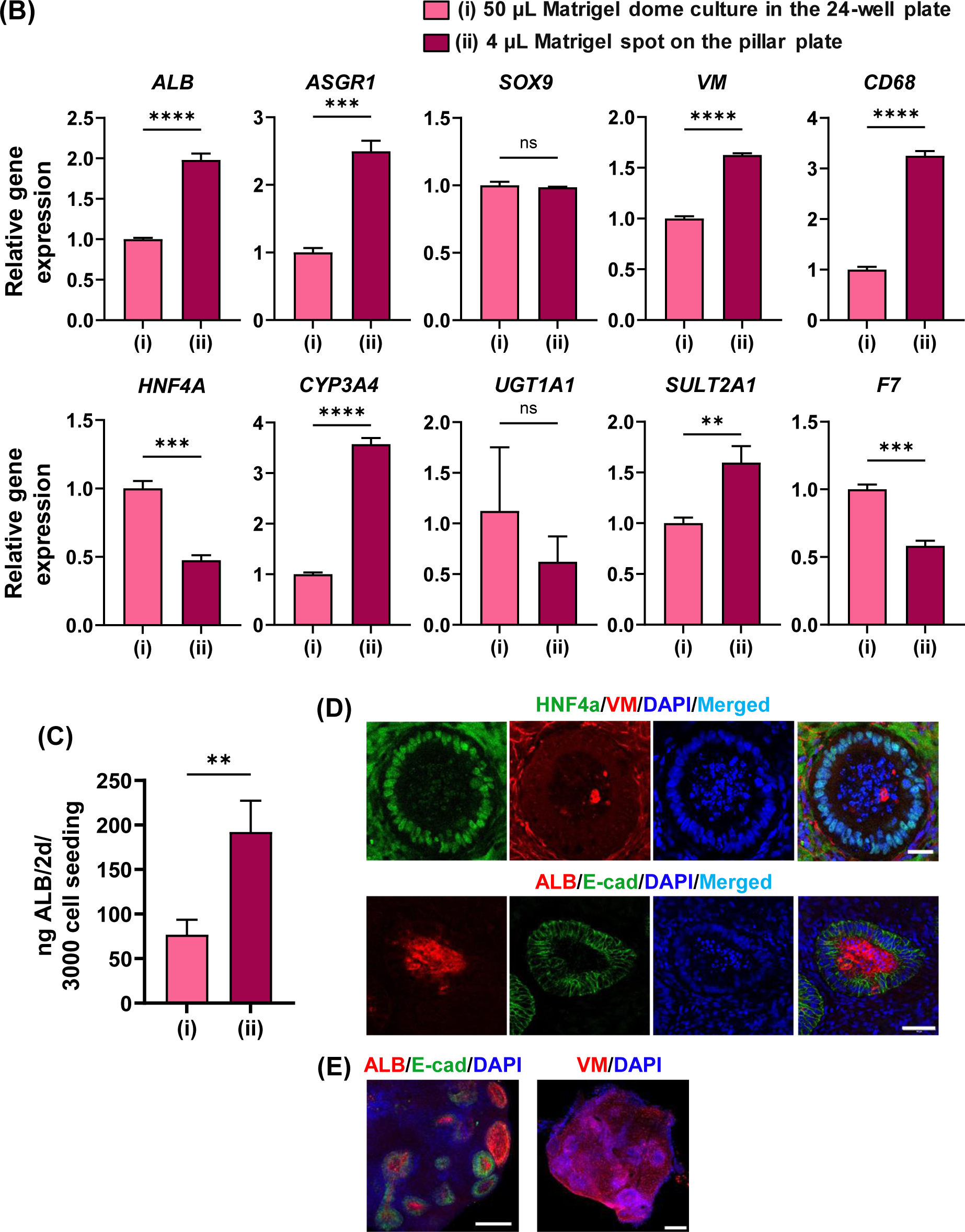
Generation of HLOs on the pillar plate. **(A)** HLOs generated in **(i)** 50 µL Matrigel dome in the 24-well plate and **(ii)** 4 µL Matrigel spot on the pillar plate. Scale bars: 200 µm except 50 µm for individual HLO. **(B)** The expression level of hepatic biomarkers including *ALB* albumin, *ASGR1* hepatocytes, *SOX9* cholangiocytes, *VM* stellate cells, *CD68* Kupffer cells, *HNF4A* early hepatic progenitor cells, drug metabolizing enzymes such as *CYP3A4*, *UGT1A1*, *SULT2A1*, and *F7* coagulation factor VII in HLOs generated **(i)** in the 24-well plate and **(ii)** on the pillar plate. N = 4 for (i) and n = 24 for (ii). **(C)** Albumin secretion measured by ELISA using HLOs generated **(i)** in the 24-well plate and **(ii)** on the pillar plate. n = 3 for (i) and n = 10 for (ii). **(D)** Immunofluorescence staining of day 25 HLOs generated on the pillar plate. Scale bars: 50 µm. **(E)** Immunofluorescence staining of whole Matrigel spot on the pillar plate. Scale bars: 200 µm.

HLOs were uniformly generated on the entire 36PillarPlate (**Figure 5A**), although the apparent shape of Matrigel spots looked different due to ECM remodeling. To assess the reproducibility of HLO generation, we first measured ATP levels of HLOs on the pillar plate using an ATP-based luminescence assay. The CV values of ATP levels in HLOs cultured on three 36PillarPlates were 16%, 15%, and 11%, respectively, with an average CV value of 14 ± 2.7%, indicating a similar number of live HLOs on the pillar plate (**Figure 5B**). The CV was used to assess the range of errors within the same trial whereas the statistical analysis was performed to measure the difference among the three trials. Since the acceptable range of CV values is 25%, HLOs generated in the same trial could be used for cell-based assays. In addition, the reproducibility of HLO function was determined by measuring the secretion of ALB and the expression of CYP3A4, a representative drug-metabolizing enzyme. The secretion levels of ALB from three trials were 152 ± 23.7, 213 ± 71.9, and 212 ± 28.0 ng per 2 days per pillar (**Figure 5C**). The average CV value calculated was 18.2%, demonstrating high reproducibility of HLO function based on ALB secretion. Furthermore, the CV value of 22% was obtained from the measurement of CYP3A4 activity (**Figure 5D**).

**Figure 5.**
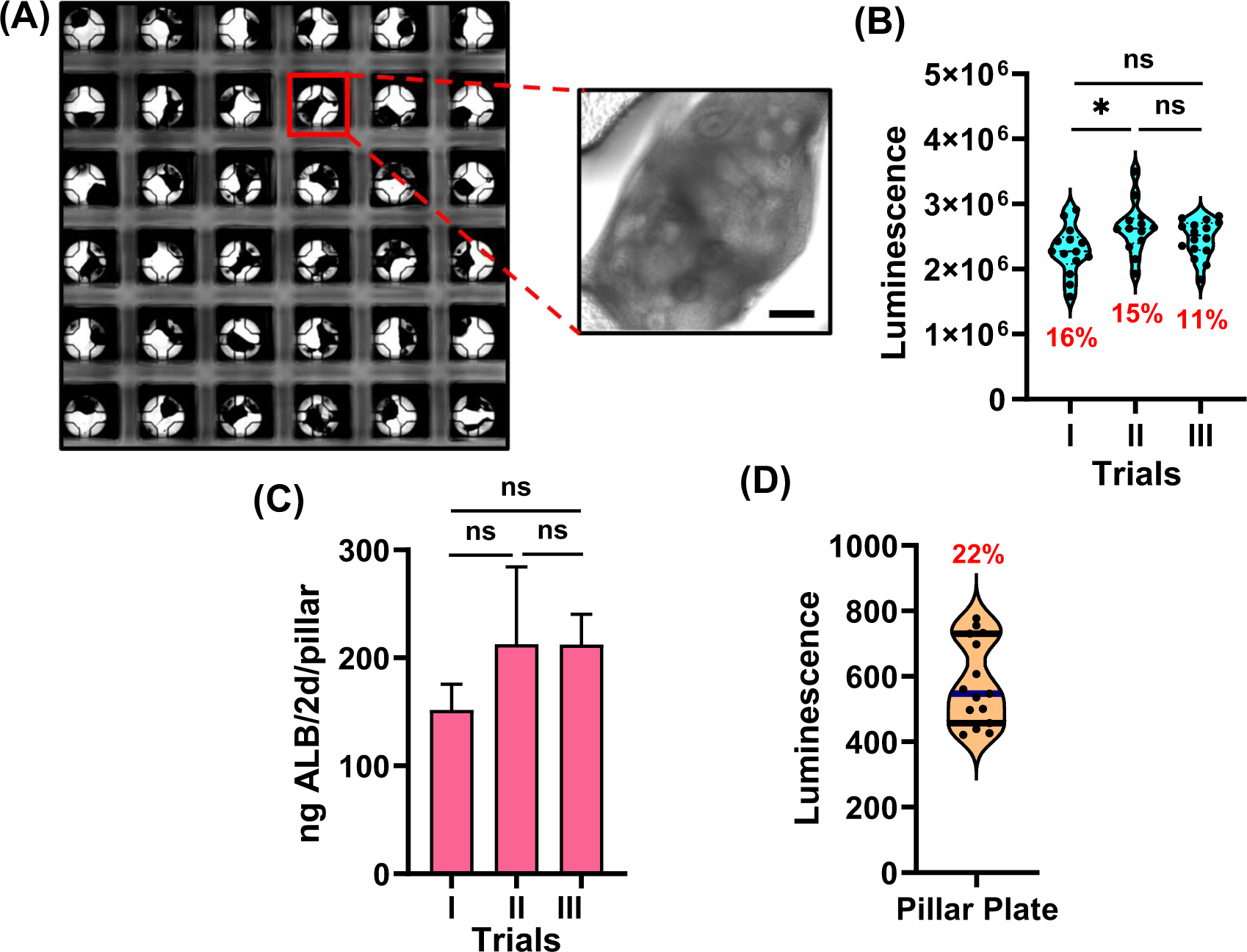
Uniform generation of HLOs on the pillar plate: **(A)** Stitched image of day 25 HLOs on the pillar plate along with a magnified image showing multiple HLOs in the Matrigel spot. Scale bar: 200 µm. **(B)** Viability of HLOs measured with the ATP-based luminescence assay. n = 16. **(C)** Albumin secretion from HLOs measured by ELISA from triplicate trials. n = 10. **(D)** CYP3A4 activity of day 25 HLOs measured by P450-Glo™ CYP3A4 luminescence assay. n = 15.

### Robustness of HLO-based in situ compound testing on the pillar plate

With the successful generation of functional HLOs on the pillar plate, we demonstrated the robustness of compound testing by generating dose-response curves of sorafenib and tamoxifen and calculating IC_50_ values in triplicate trials (**Figure 6**). Briefly, the pillar plate with HLOs was sandwiched onto the deep well plate with the two drugs (6 concentrations and 6 replicates). After drug treatment for 3 days, the viability of HLOs was measured with the ATP-based luminescence assay to generate dose-response curves. The IC_50_ values of sorafenib from triplicate trials were 6.0, 4.7, and 8.0 µM, with an average IC_50_ value of 6.2 ± 1.6 µM (**Figure 6B**). Here, a trial means an independently performed culture for the generation of day 25 HLOs on the pillar plate starting from the different batches of iPSCs. Notably, the IC_50_ values of sorafenib obtained from HLOs on the pillar plate closely aligned with previously reported IC_50_ values: 5.3 - 8.5 µM from HepG2 cells, 4.7 - 17.1 µM from Huh7 cells, and 3.3 µM from Hep3B cells ^41–44^. Sorafenib, a multi-kinase inhibitor used in the therapy of renal, liver, and thyroid cancers ^45^, is susceptible to liver injury through the production of toxic metabolites, and drug-drug interactions. It inhibits or induces hepatic CYP3A4 activity and inhibits UGT activity, resulting in hyperbilirubinemia ^45^. It is metabolized mainly by CYP3A4 and UGT1A9 ^46,47^.

**Figure 6.**
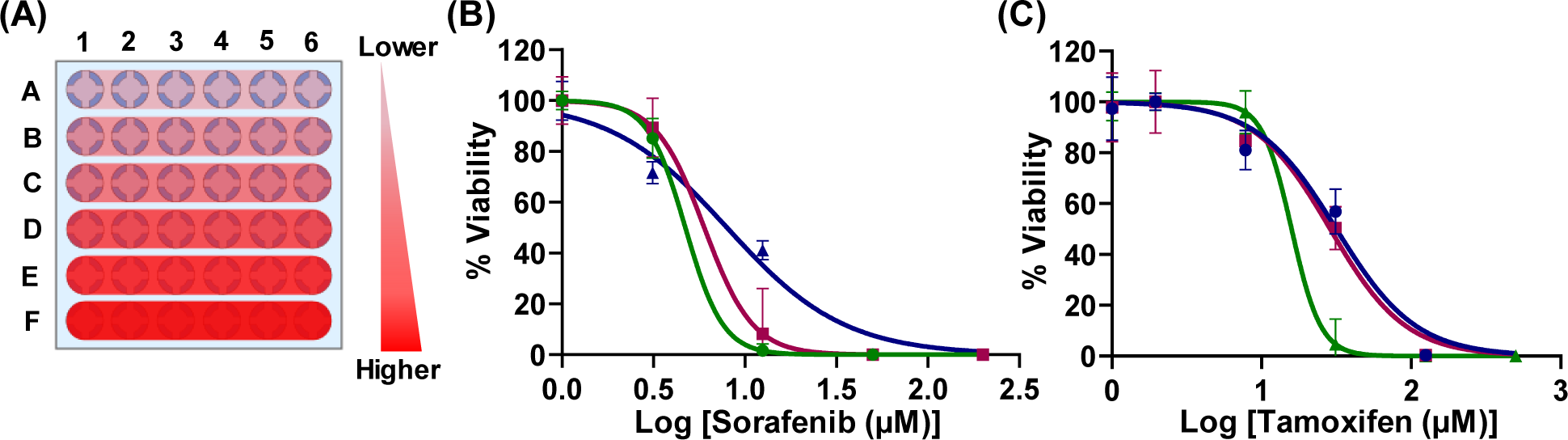
Robustness of compound testing on the pillar plate. **(A)** The dose of a compound on the pillar plate (6 concentrations and 6 replicates). **(B)** Dose response curves of sorafenib from triplicate trials. An average IC_50_ value of 6.2 ± 1.6 µM was obtained. n = 6 per each concentration. **(C)** Dose response curves of tamoxifen from triplicate trials. An average IC_50_ value of 25.4 ± 8.3 µM was obtained. n = 6 per each concentration.

For tamoxifen, the IC_50_ values obtained from three independent trials were 31.6, 15.9, and 28.8 µM, with an average IC_50_ value of 25.4 ± 8.3 µM (**Figure 6C**). The IC_50_ values for tamoxifen reported previously using HepG2 cells were 2.3 - 5.3 µM ^48,49^, considerably lower than those obtained from HLOs on the pillar plate. However, the IC_50_ value for non-malignant liver cells was 33 µM ^48^, demonstrating similarity to our findings. Additionally, the IC_50_ values of tamoxifen from 2D and 3D cultured iCell hepatocytes (iPSC-derived hepatocytes) were 10.2 and 12.4 µM, respectively, whereas the IC_50_ value obtained from HepG2 cell spheroids was 18.7 ± 29.4 µM ^50^, which are also in line with our findings. Tamoxifen, a nonsteroidal antiestrogen widely used in the treatment of breast cancer ^51^, is hepatotoxic and induces fatty liver, steatohepatitis, cirrhosis, and acute liver injury ^51^. It is metabolized mainly by phase I enzymes such as CYPs and FMOs and phase II enzymes such as SULTs and UGTs ^52^.

## Discussion

For high-throughput, organoid-based compound screening, it is critical to generate mature organoids reproducibly and robustly in a miniature system that can accommodate *in situ* organoid imaging and functional testing. Conventional organoid culture systems including 6-/24-well plates with Matrigel domes, micropatterned well plates, petri dishes, and spinner flasks require relatively large volume of expensive organoid culture media and are low throughput due to incompatibility with common analytical equipment ^53^. For example, *in situ* imaging of organoids in Matrigel domes in 6-/24-well plates could be challenging due to the thickness of Matrigel domes, and large Matrigel domes could induce diffusion limitation of nutrients and oxygen in the core, leading to variability in organoid differentiation ^54^. Thus, it was necessary to isolate and transfer harvested organoids from conventional organoid culture systems to a high-density well plate for compound screening. For instance, HLOs cultured in Matrigel domes in the 24-well plate were harvested by mechanical scratching and pipetting and dispensed in an ultralow attachment (ULA) 384-well plate for high-throughput, drug screening assays ^12^. The major concern of this approach is that the mechanical/enzymatic dissociation of organoids in Matrigel domes is labor-intensive and could lead to the loss of organoids, damage to organoid structure and function, and batch-to-batch variation in their size.

To avoid these issues and improve organoid homogeneity, hydrogel-coated microcavity plates have been developed for high-throughput, static culture of human gastrointestinal organoids ^55^. This approach is a significant advancement in the field of organoid research. While it is ideally suitable for the short-term differentiation of organoids, it could not support dynamic organoid culture for high maturity and differentiation of progenitor cells that inevitably require hydrogel encapsulation. Although miniature organoid culture in microfluidic devices looks promising, it is difficult to load cells rapidly in microchannels and microchambers and upscale organoid generation ^56^. In this study, we addressed these issues by developing the pillar plate with sidewalls and slits and demonstrated rapid 3D bioprinting of foregut cells in Matrigel and differentiation of HLOs on the pillar plate for *in situ* organoid imaging and hepatotoxicity testing. The pillar plate coupled with the deep well plate streamlines the process of organoid differentiation and compound screening by enabling rapid loading of cells on the pillar plate with minimal manual intervention and long-term culture of spheroids/organoids in hydrogels with simple culture medium change by sandwiching. Since the pillar plate is built on the footprint of standard 384-well plates, common lab equipment such as microtiter well plate readers and automated fluorescence microscopes can be used to acquire absorbance, fluorescence, and luminescence signals from organoids directly without the harvesting steps, which is critical for high-throughput, organoid-based assays. Furthermore, the pillar plate miniaturizes organoid culture, requiring only 4 - 5 μL of cells in hydrogels on each pillar (200 – 3,000 cells/pillar) and 80 μL of culture media in each deep well. Thus, it reduces the resources by 12 - 100-fold as compared to conventional 6-/24-well plates, petri dishes, and spinner flasks. Finally, statically cultured organoids on the pillar plate can be exposed to dynamic environments at any time points by simply sandwiching the pillar plate onto a perfusion plate ^57,58^, or dynamically cultured organoids in the pillar/perfusion plate can be tested with compounds by simply sandwiching the pillar plate onto the deep well plate. This kind of flexibility has not been demonstrated by any existing organoid culture platforms.

In this study, we demonstrated high-throughput, miniature foregut cell printing in Matrigel on the pillar plate by using a microsolenoid valve-driven 3D bioprinter, generated day 25 HLOs uniformly, and performed *in situ* assessment of compound hepatotoxicity. Microsolenoid valve-driven, microarray 3D bioprinting is ideally suited for dispensing fragile cells, including iPSCs and organoids, and could offer high printing accuracy and speed, low shear stress, and high viability (< 98%) for bioprinted cells ^59,60^. The printing volume in the range of 100 nL – 6 µL can be varied easily by changing the valve open time and pressure. While achieving high viability and uniform printing of cells on the pillar plate (**Figure 2**), the printing speed was extremely fast: 20 and 60 seconds for the 36PillarPlate and the 384PillarPlate, respectively. In addition, microsolenoid valve-driven, microarray 3D bioprinting can accommodate various hydrogels including Matrigel, BME2, Geltrex, collagen, and alginate (data not shown). Since Matrigel has been used as the gold standard for organoid differentiation and maturation ^61–64^, we generated HLOs in Matrigel spots on the pillar plate. Due to mild thermal crosslinking, bioprinted iPSCs and foregut cells in Matrigel were highly viable (**Figure 2**).

Recently, our group demonstrated the proof of concept of organoid printing on the pillar plate by using harvested HLOs from Matrigel domes for studying steatohepatitis ^57^. In addition, we demonstrated uniform cell spheroid transfer to the pillar plate for robust and reproducible culture of human brain organoids ^65^. Thus, in the present work, *in situ* generation of HLOs on the pillar plate was demonstrated by printing foregut cells in Matrigel and directly differentiating them on the pillar plate for hepatotoxicity testing. Using commercially available iPSCs, we performed optimization of bioprinting protocols and provided the optimum conditions to demonstrate high viability of printed iPSCs and foregut cells in Matrigel and high bioprinting accuracy. Printing parameters were optimized with a more advanced 3D bioprinter, which uses disposable printing tips without rinsing and drying steps, leading to extremely fast cell printing. In addition, in-depth comparison of HLOs generated on the pillar plate and in the 24-well plate (i.e., conventional Matrigel dome culture), including immunofluorescence staining, hepatic gene expression, and other functional assays, were performed. The reproducibility of HLO generation on the pillar plate has been demonstrated by showing the stitched image of the entire 36PillarPlate and measuring ATP and albumin levels in three independent trials (**Figure 5**). In addition, the robustness of *in situ* compound testing has been demonstrated with HLOs on the pillar plate in three independent trials (**Figure 6**). Thus, we envision that the pillar plate coupled with microarray 3D bioprinting technology could be utilized in industrial and clinical settings for high-throughput, *in situ* assessment of compounds with organoids.

Although HLOs represent a promising tool for liver disease modeling and predictive hepatotoxicity screening, it is still challenging to replace primary human hepatocytes with HLOs due to their limited maturity, particularly in drug metabolism. With the most advanced HLO differentiation protocol currently available (**Figure 3A**), the overall expression level of drug metabolizing enzymes in HLOs is 10- to 1000-fold lower than that in primary human hepatocytes ^6,12^. Thus, HLOs have been used mainly for recapitulating a specific function of the liver and diseases, including steatohepatitis ^6^ and drug-induced cholestatic liver injury ^12^. With limited maturity of HLOs compared to primary hepatocytes, it would be still challenging to test metabolism-sensitive compounds such as acetaminophen accurately. Significant efforts should be devoted to enhancing the maturity of human organoids.

## Conclusions

We have successfully demonstrated the robust and reproducible generation of liver organoids by printing foregut cells in Matrigel on the pillar plate and differentiating them in the deep well plate. The microsolenoid value-driven, microarray 3D bioprinting technology allowed us to print fragile iPSCs and foregut cells in Matrigel on the pillar plate uniformly and achieve exceptionally high cell viability. The printing speed was extremely fast as compared to conventional 3D bioprinting technologies, which could be important for maintaining high cell/organoid viability after printing. The pillar plate supported miniature organoid culture with at least 12-folds and up to 100-folds reduced medium volume as compared to traditional organoid culture systems. In addition, we demonstrated the feasibility of *in situ* hepatotoxicity assessment on the pillar plate by exposing HLOs to compounds and fluorescent/luminescent reagents without transferring organoids to microtiter well plates for biological assays. Therefore, by automating and miniaturizing the organoid culture system by using the pillar plate and microarray 3D bioprinting technology, we have successfully generated reproducible liver organoids that are compatible for *in situ* assessments. This capability could be critical for organoid-based, high-throughput compound screening.

## Abbreviations

36-pillar plate: with sidewalls and slits
36PillarPlate: 384-pillar plate with sidewalls and slits
384PillarPlate: clear bottom 384-deep well plate
384DeepWellPlate: induced pluripotent stem cells
iPSCs: human liver organoid
HLO: high-throughput screening
HTS: coefficient of variation
CV: vimentin
VM: albumin
ALB: hepatocyte nuclear factor 4 alpha
HNF4a: SRY-box transcription factor 9
SOX9: E-cadherin
E-cad: asialoglycoprotein receptor 1
ASGR1: cluster of differentiation 68
CD68: cytochrome P450
CYP450: uridine-5’-diphospho-glucuronosyl transferases
UGT: flavin monooxygenase
FMO: sulfotransferase
SULT: coagulation factor VII F7

## Acknowledgments

This study was financially supported by the National Institutes of Health (NIDDK UH3DK119982, NCATS R44TR003491, and NIEHS R43ES035653) and the Ohio Third Frontier Commission (TVSF Phase IB and Phase II).

---

**Supplementary Figure 1.**
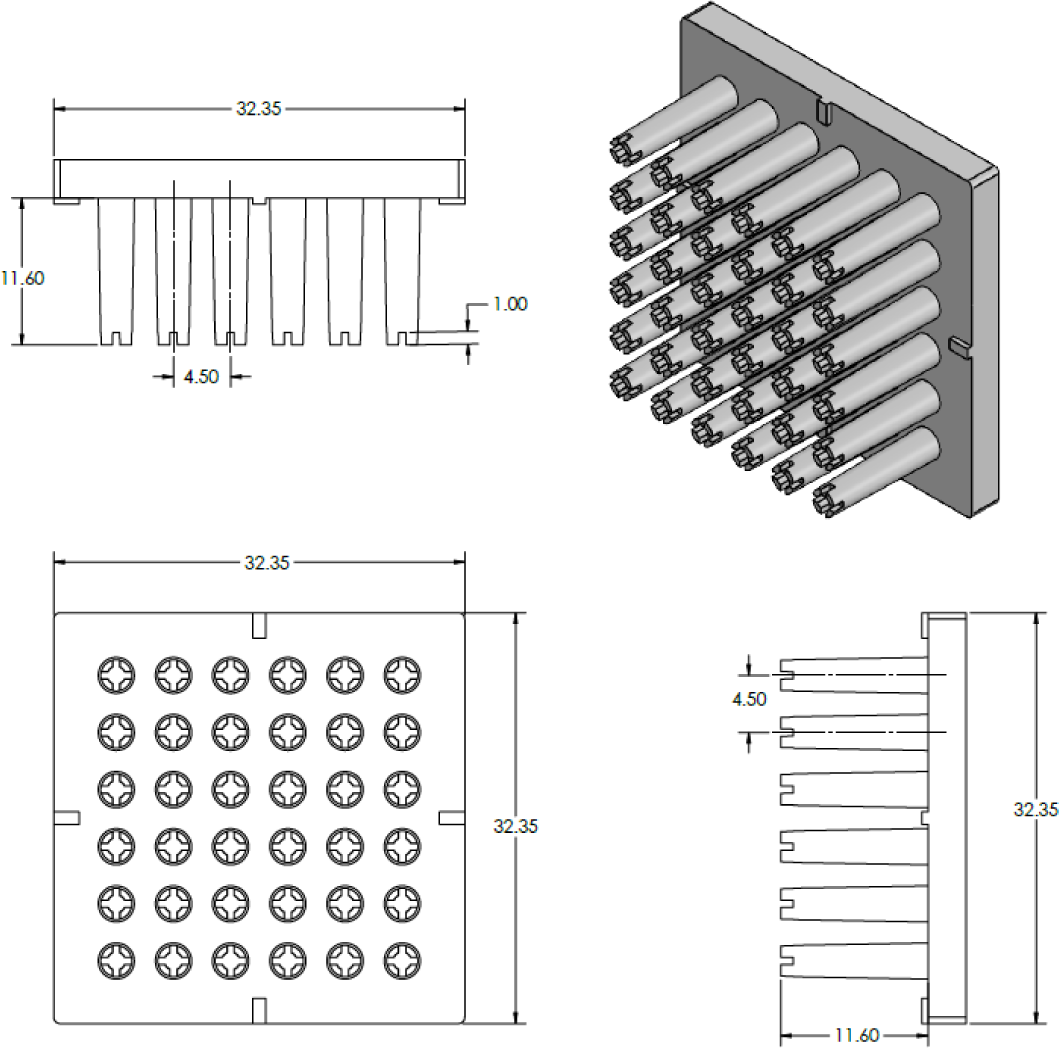
SolidWorks design of the 36PillarPlate with a 6 x 6 array of pillars. The unit of the dimension is millimeter (mm).

**Supplementary Figure 2.**
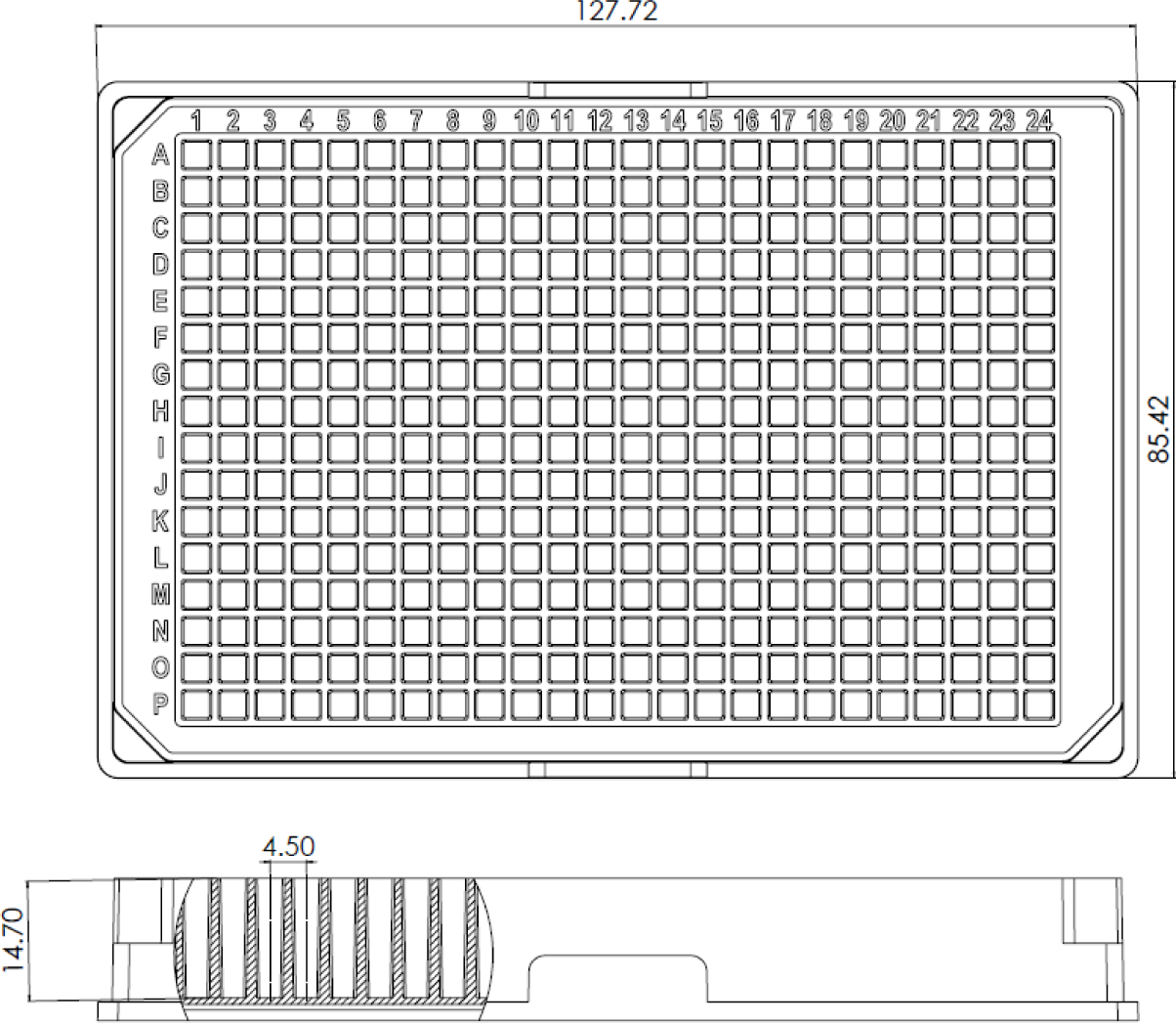
SolidWorks design of the 384DeepWellPlate with a 16 x 24 array of deep wells. The unit of the dimension is millimeter (mm).

**Supplementary Figure 3.**
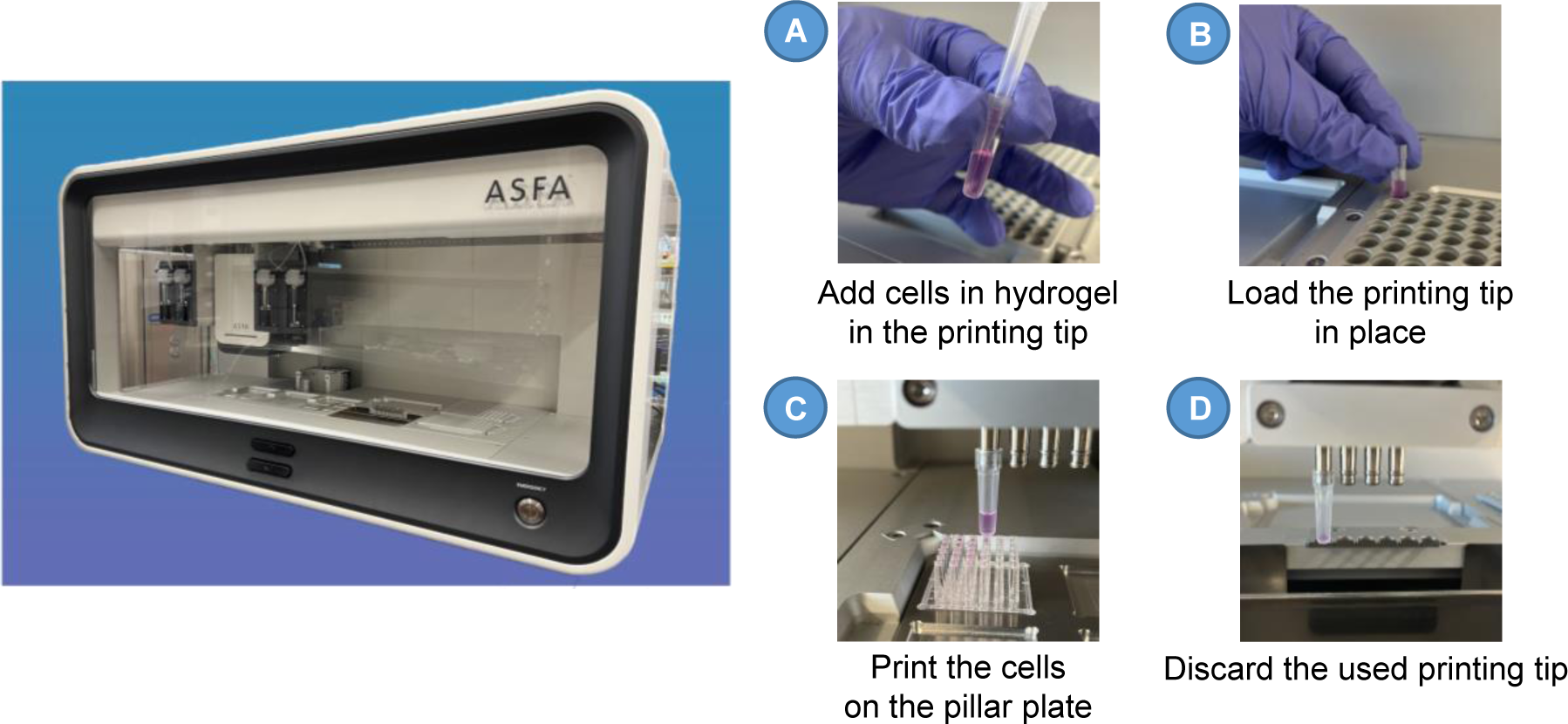
The 3D bioprinter used for cell printing on the pillar plate. The 3D bioprinter is operated by microsolenoid valves and pneumatic pressure for sample printing. The pictures A – D illustrate crucial steps necessary for printing cells in hydrogel on the pillar plate.

**Supplementary Figure 4.**
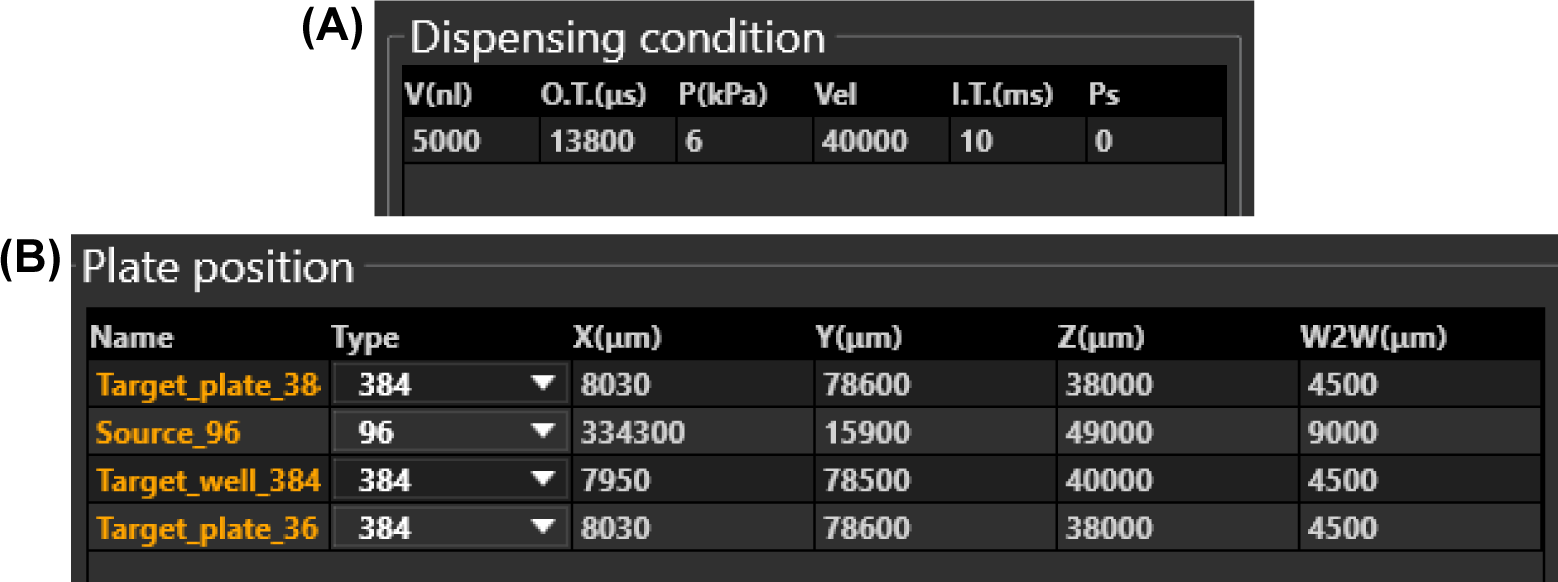
**(A)** Example of a sample dispensing condition. To dispense 5 µL of samples, 13,800 µs of open time (O.T.) and 6 kPa of pressure (P) were used. **(B)** The X, Y, and Z position and the well-to-well (W2W) or pillar-to-pillar distance of sample plates and pillar plates in µm.

**Supplementary Figure 5.**
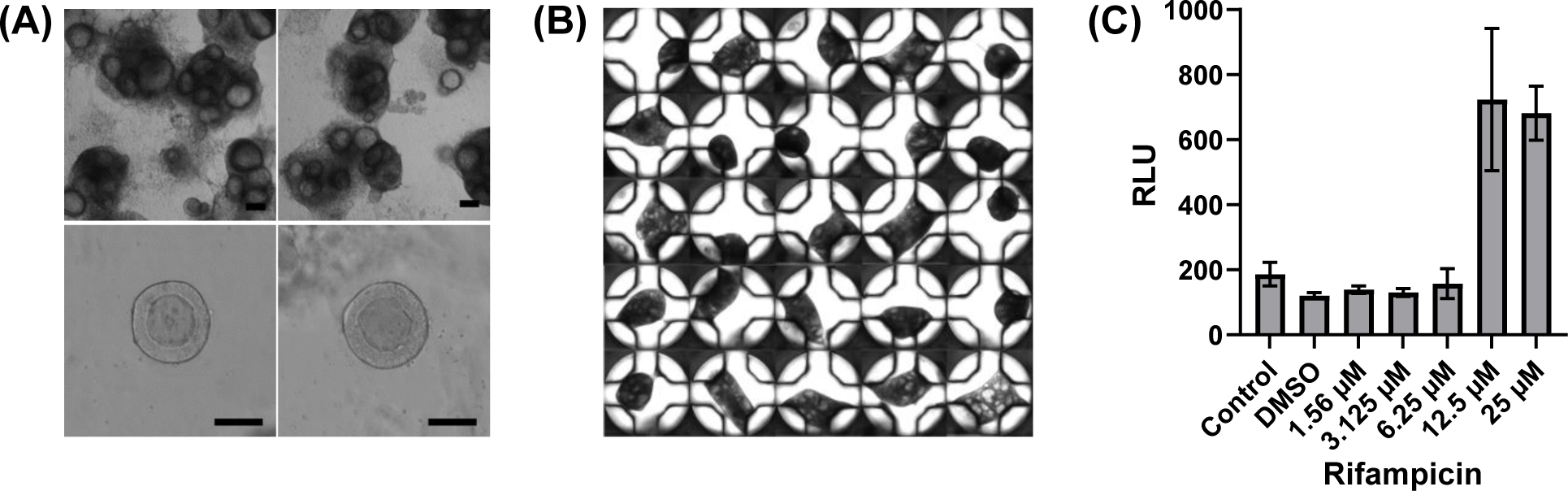
Generation of HLOs by differentiation of 72-3 iPSCs. **(A)** Representative images of HLOs generated in 50 µL Matrigel domes in a 24-well plate (Upper row, scale bars: 200 µm) and magnified images of individual HLOs (Bottom row, scale bars: 50 µm). Stitched image of day 25 HLOs on the 384PillarPlate. Day 7 foregut cells were suspended in 2-fold diluted Matrigel and printed on the pillar plate at the seeding density of 3,000 cells/pillar. Induction of CYP3A4 by treatment of HLOs with rifampicin at the concentration range of 1.6 µM - 25 µM. The control condition contains no DMSO and no rifampicin whereas the DMSO condition contains 0.5% DMSO alone in the culture medium. All rifampicin treatment conditions contain 0.5% DMSO.

**Supplementary Table 1.**
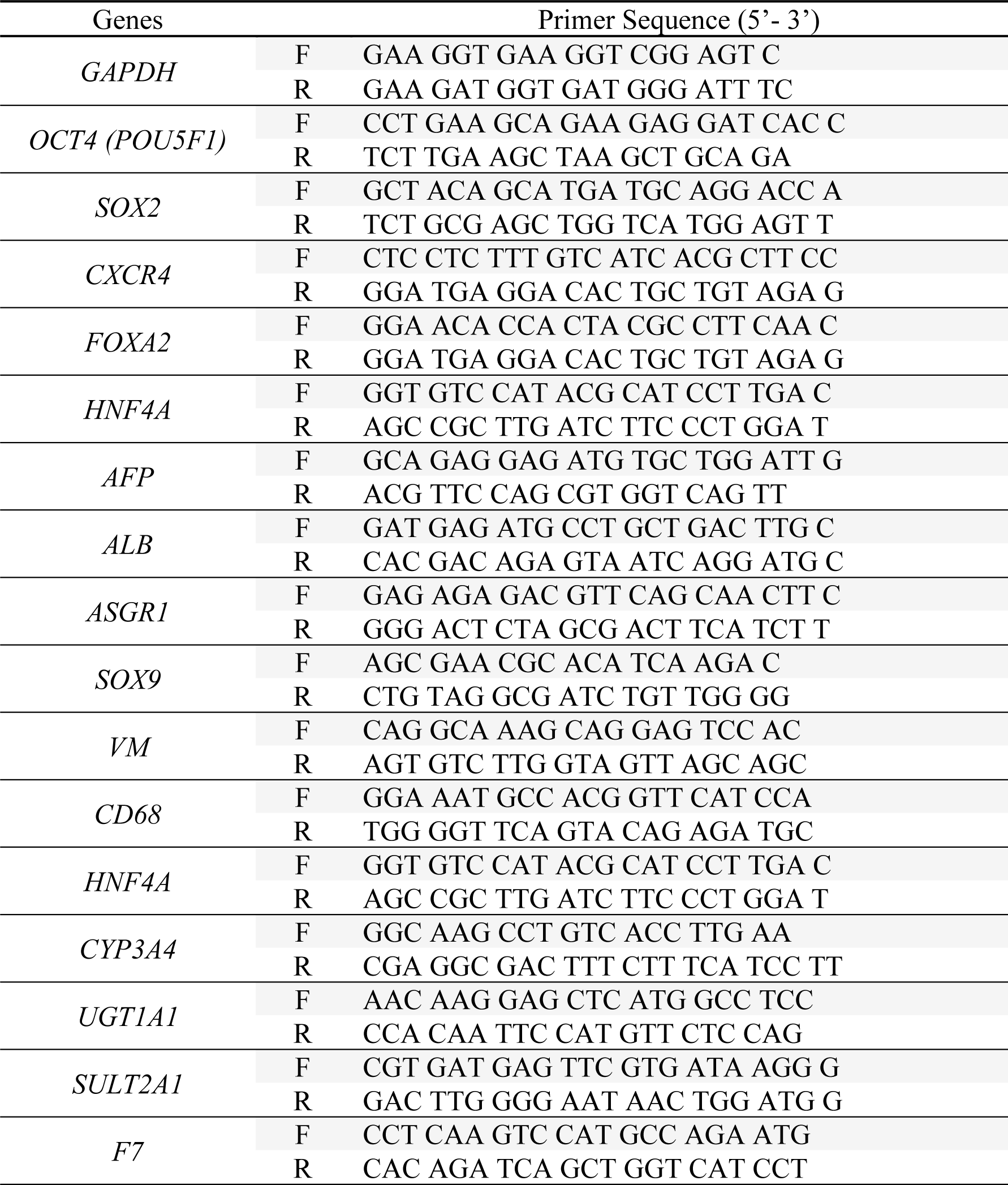
List of primers.

**Supplementary Table 2.**
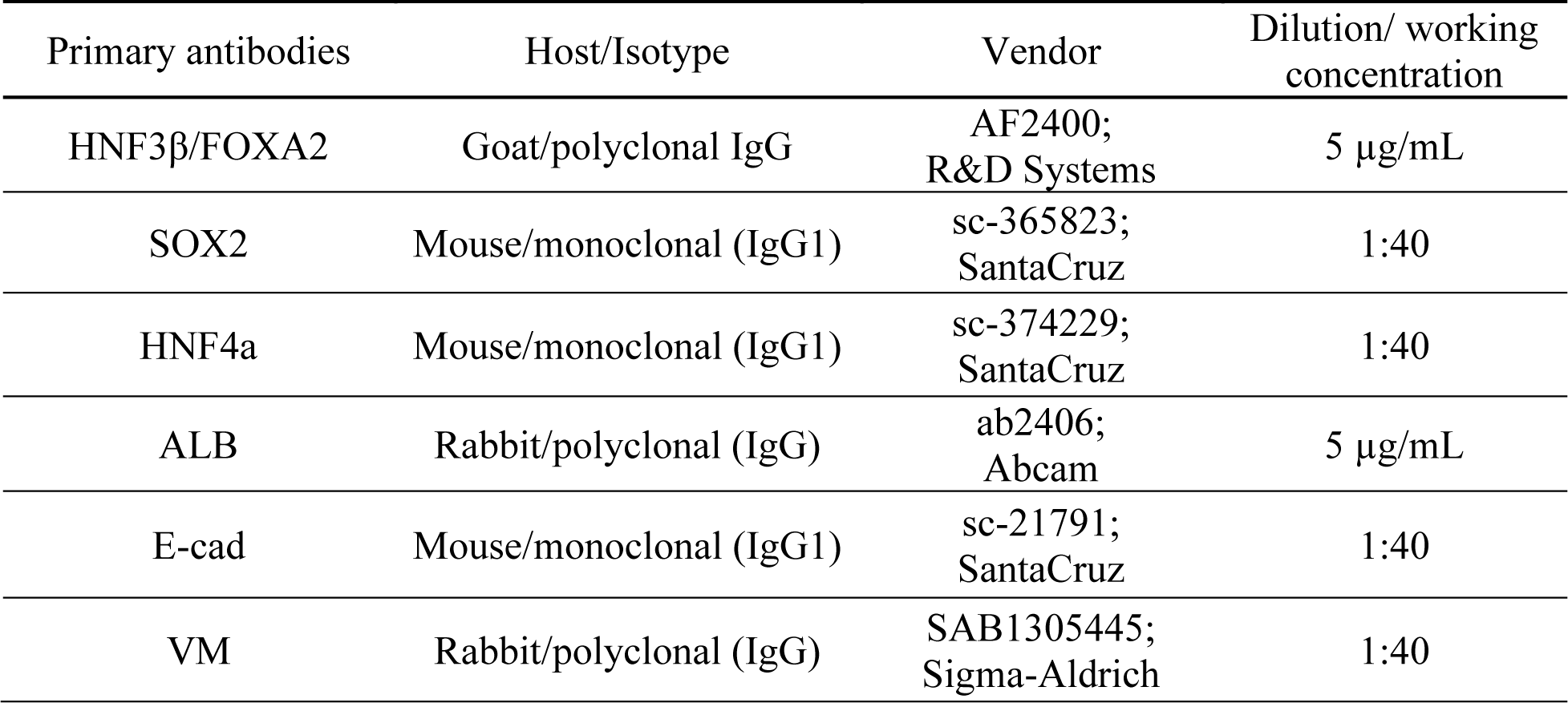
List of Primary antibodies.

| Primary antibodies | Host/Isotype | Vendor | Dilution/ working concentration |
| --- | --- | --- | --- |
| HNF3β/FOXA2 | Goat/polyclonal IgG | AF2400;<br>R&D Systems | 5 μg/mL |
| SOX2 | Mouse/monoclonal (IgG1) | sc-365823;<br>SantaCruz | 1:40 |
| HNF4a | Mouse/monoclonal (IgG1) | sc-374229;<br>SantaCruz | 1:40 |
| ALB | Rabbit/polyclonal (IgG) | ab2406;<br>Abcam | 5 μg/mL |
| E-cad | Mouse/monoclonal (IgG1) | sc-21791;<br>SantaCruz | 1:40 |
| VM | Rabbit/polyclonal (IgG) | SAB1305445;<br>Sigma-Aldrich | 1:40 |

**Supplementary Table 3.** List of Secondary antibodies.

| Secondary antibodies | Host | Vendor | Dilution |
| --- | --- | --- | --- |
| Donkey anti-Mouse IgG (H+L) Highly Cross-Adsorbed Secondary Antibody, Alexa Fluor™ 488 | Donkey | A-21202;<br>Invitrogen | 1:400 |
| Donkey anti-Mouse IgG (H+L) Highly Cross-Adsorbed Secondary Antibody, Alexa Fluor™ 647 | Donkey | A-31571;<br>Invitrogen | 1:400 |
| Donkey anti-Rabbit IgG (H+L) Highly Cross-Adsorbed Secondary Antibody, Alexa Fluor 488 | Donkey | A-21206;<br>Invitrogen | 1:400 |
| Donkey anti-Rabbit IgG (H+L) Highly Cross-Adsorbed Secondary Antibody, Alexa Fluor 647 | Donkey | A-31573;<br>Invitrogen | 1:400 |
| Donkey anti-Goat IgG (H+L) Highly Cross-Adsorbed Secondary Antibody, Alexa Fluor 647 | Donkey | A-21447;<br>Invitrogen | 1:400 |

## Notes

### Competing Interest Statement

The authors have declared no competing interest.

